# *Escherichia ruysiae* sp. nov., isolated from an international traveller

**DOI:** 10.1101/781724

**Authors:** Boas C.L. van der Putten, S. Matamoros, D.R. Mende, COMBAT consortium, C. Schultsz

## Abstract

The *Escherichia* genus comprises four species and at least five lineages currently not assigned to any species, termed ‘*Escherichia* cryptic clades’. We isolated an *Escherichia* strain from an international traveller and resolved the complete DNA sequence of the chromosome and an IncI multi-drug resistance plasmid using Illumina and Nanopore whole-genome sequencing (WGS). Strain OPT1704^T^ can be differentiated from existing *Escherichia* spp. using biochemical (VITEK2) and genomic tests (average nucleotide identity [ANI] and digital DNA:DNA hybridisation [dDDH]). Phylogenetic analysis based on alignment of 16S rRNA sequences and 682 concatenated core genes showed similar results. Our analysis further revealed that strain OPT1704^T^ falls within *Escherichia* cryptic clade IV, and is closely related to cryptic clade III. Combining our analyses with publicly available WGS data of cryptic clades III and IV from Enterobase confirmed the close relationship between clades III and IV (>96% interclade ANI), warranting assignment of both clades to the same novel species. We propose *E. ruysiae* sp. nov. as a novel species, encompassing *Escherichia* cryptic clades III and IV (type strain OPT1704^T^ = NCCB 100732^T^ = NCTC 14359^T^).

**Author notes:** The Genbank accession number for the 16S rRNA gene sequence of strain OPT1704^T^ is LR745848. The Genbank accession number for the complete genome sequence of strain OPT1704^T^ is CABVLQ000000000.

## Introduction

Within the *Escherichia* genus, four species are recognized; *E. coli* (1), *E. fergusonii* (2), *E. albertii* (3) and most recently, *E. marmotae* (4). Several species were assigned to the *Escherichia* genus previously, but have now been moved to other genera, such as *E. vulneris* (now *Pseudescherichia vulneris* (5)), *E. blattae* (now *Shimwellia blattae* (6)), *E. adecarboxylata* (now *Leclercia adecarboxylata* (7)) and *E. hermannii* (now *Atlantibacter hermannii* (8)). All four *Escherichia* species have been associated with the potential to cause animal and/or human disease (9–12). Several *Escherichia* strains cannot be assigned to any of the four existing species. Based on analysis of genetic data, these strains cluster into several groups, which were termed ‘*Escherichia* cryptic clades’ (13,14). Recently, cryptic clade V was formally recognized as a separate species (*E. marmotae*), leaving five cryptic clades that have not been delineated at the species level. Here we report the novel species *Escherichia ruysiae* sp. nov., isolated from faecal material of an international traveller. *Escherichia ruysiae* sp. nov. encompasses the closely related *Escherichia* cryptic clades III and IV.

## Isolation and Ecology

We discovered a cryptic clade IV strain in our collection, previously identified as extended spectrum beta-lactamase (ESBL) producing *E. coli* as part of the COMBAT study, which investigated the acquisition of ESBL-producing Enterobacteriaceae (ESBL-E) during international travel (15). This isolate, OPT1704^T^, was further characterized in detail.

The strain was isolated from a human faecal sample provided immediately after an individual’s return from a one-month journey to several Asian countries. No ESBL-E were detected in a faecal sample collected immediately before departure, suggesting the ESBL gene, and possibly strain OPT1704^T^, were acquired during travel. The traveller reported diarrhoea during travel but no antibiotic usage. No ESBL-E were isolated in follow-up faecal samples, suggesting loss of the OPT1704^T^ strain or the ESBL gene within one month after return from travel.

## Genome Features

The whole-genome sequence of strain OPT1704^T^ was determined using a combination of the Illumina HiSeq and Oxford Nanopore Technologies (ONT) sequencing platforms. The Illumina sequencing run yielded a total of 6.3×10^6^ paired-end reads, with a mean read length of 151 bp. Illumina reads were downsampled using seqtk (version 1.3-r106, https://github.com/lh3/seqtk) to provide a theoretical coverage depth of 100X with the assumption that the OPT1704^T^ has a genome size of approximately 5×10^6^ bp. The ONT sequencing run yielded a total of 2.5×10^4^ reads, with a mean read length of 9078 bp before filtering. ONT reads were filtered on length and on read identity using Filtlong (version 0.2.0, https://github.com/rrwick/Filtlong) with Illumina reads as a reference, leaving 1.5×10^4^ reads with a mean length of 12580 bp. This provided a theoretical coverage depth of ∼38X of ONT reads. The combined assembly using Unicycler (version 0.4.6 (16)) of Illumina and Nanopore reads resulted in a completely assembled genome, consisting of one circular chromosome and one circular plasmid. The GC content of the complete OPT1704^T^ genome was 50.6%.

Putative resistance and virulence genes were predicted from the draft genome using ABRicate with the CARD (17) and VFDB (18) databases. OPT1704^T^ harbours 6 resistance genes on its IncI plasmid, associated with reduced susceptibility to fluoroquinolones (*qnrS1*), aminoglycosides (*aph(6)-Id* & *aph(3’’)-Ib*), cephalosporins (*blaCTX-M-14*), trimethoprim (*dfrA14*) and sulphonamides (*sul2*), corresponding with its reduced susceptibility to fluoroquinolones (norfloxacin, MIC: 2 mg/L and ciprofloxacin, MIC: 0.5 mg/L), cephalosporins (cefuroxime, MIC: >32 mg/L and cefotaxime, MIC: 4 mg/L) and trimethoprim-sulfamethoxazole (MIC: >8 mg/L), assessed using VITEK2 (BioMérieux). However, strain OPT1704T was susceptible to tobramycin (MIC: ≤1 mg/L) and gentamicin (MIC: ≤1 mg/L) despite presence of aminoglycoside resistance genes. Furthermore, several putative virulence genes were predicted from the genome sequence associated with siderophore function (*chuX, entS, fepABD*), fimbriae (*fimBCDGI*), a type II secretion system (*gspGHI*) and capsular polysaccharide biogenesis (*kpsD*). These predicted virulence genes, when present in *Escherichia coli*, are not typically associated with a specific clinical syndrome such as diarrhoeal disease.

## Physiology and Chemotaxonomy

Strain OPT1704^T^ formed circular, grey-white colonies on a Columbia sheep (COS) blood agar plate. Individual cells were observed under a light microscope and were rod-shaped and approximately 1 by 2 µm in size. The strain was shown to be Gram-negative, non-motile, oxidase-negative and catalase-positive. The strain was capable to grow in the absence of oxygen. On COS blood plates, it showed growth in the temperature range of 20-42 °C. The strain was also able to grow in NaCl concentrations ranging from 0% to 6% in lysogeny broth. MALDI-TOF (Bruker) and VITEK2 (BioMérieux) systems both identified OPT1704^T^ as *E. coli* with high confidence scores (score>2 for MALDI-TOF and “Excellent identification” for VITEK2). Comparison of the output of the VITEK2 biochemical test with published biochemical reactions of other *Escherichia* species revealed that *E. ruysiae* sp. nov. str. OPT1704^T^ is distinct from other *Escherichia* species based on a biochemical profile (table 1) (2–4,19). One biochemical reaction, lysine decarboxylation, cannot be performed by strain OPT1704^T^, but can be performed by all other *Escherichia* species. This reaction is typically mediated by the cadA gene (20), which is missing in OPT1704^T^ but present in other *Escherichia*.

**Table 1.** Comparison of biochemical markers which differentiate *E. ruysiae* sp. nov. from other *Escherichia* species. + and – indicate that ≥85% of tested strains is positive or negative for that biochemical marker, respectively. Data for *E. albertii, E. coli, E. fergusonii* and *E. marmotae* summarised from literature (Abbott 2003, Huys 2003, Farmer 1985, Liu 2015).

|  | *E. ruysiae* | *E. albertii* | *E. coli* | *E. fergusonii* | *E. marmotae* |
| --- | --- | --- | --- | --- | --- |
| ONPG | + | + | + | + | ? |
| Lysine decarboxylase | - | + | + | + | + |
| Ornithine decarboxylase | + | + | + | + | ? |
| Fermentation of: |  |  |  |  |  |
| Adonitol | - | ? | ? | + | ? |
| d-Xylose | - | ? | + | + | + |
| Cellobiose | - | ? | ? | + | ? |
| d-Sorbitol | + | ? | + | ? | + |

\*50-85% of *E. coli* possess this biochemical property.

## 16S rRNA and whole-genome phylogeny

Next, we calculated 16S rRNA sequence similarities, ANI values and digital DNA:DNA hybridisation (dDDH) values between OPT1704^T^ and type strains of the four other *Escherichia* species, representative genomes of the other three *Escherichia* cryptic clades, and *S. enterica* serovar Typhimurium (table 2). Representative genomes for the *Escherichia* cryptic clades I, II, III and VI were selected from Enterobase (21), using the genomes with the highest contiguity. Clades VII and VIII in Enterobase only consisted of a single strain and were not used in further analyses. We used three separate tools to calculate average nucleotide identity (ANI) (fastANI (22), OrthoANIu (23) and ANI calculator from Enveomics (24)) and the DSMZ Genome-to-Genome Distance Calculator to calculate digital DNA:DNA hybrisation (dDDH) (25). 16S rRNA genes were extracted from whole genomes using barrnap (version 0.9, https://github.com/tseemann/barrnap) and similarity was assessed using snp-dists (version 0.6, https://github.com/tseemann/snp-dists).

**Table 2.**
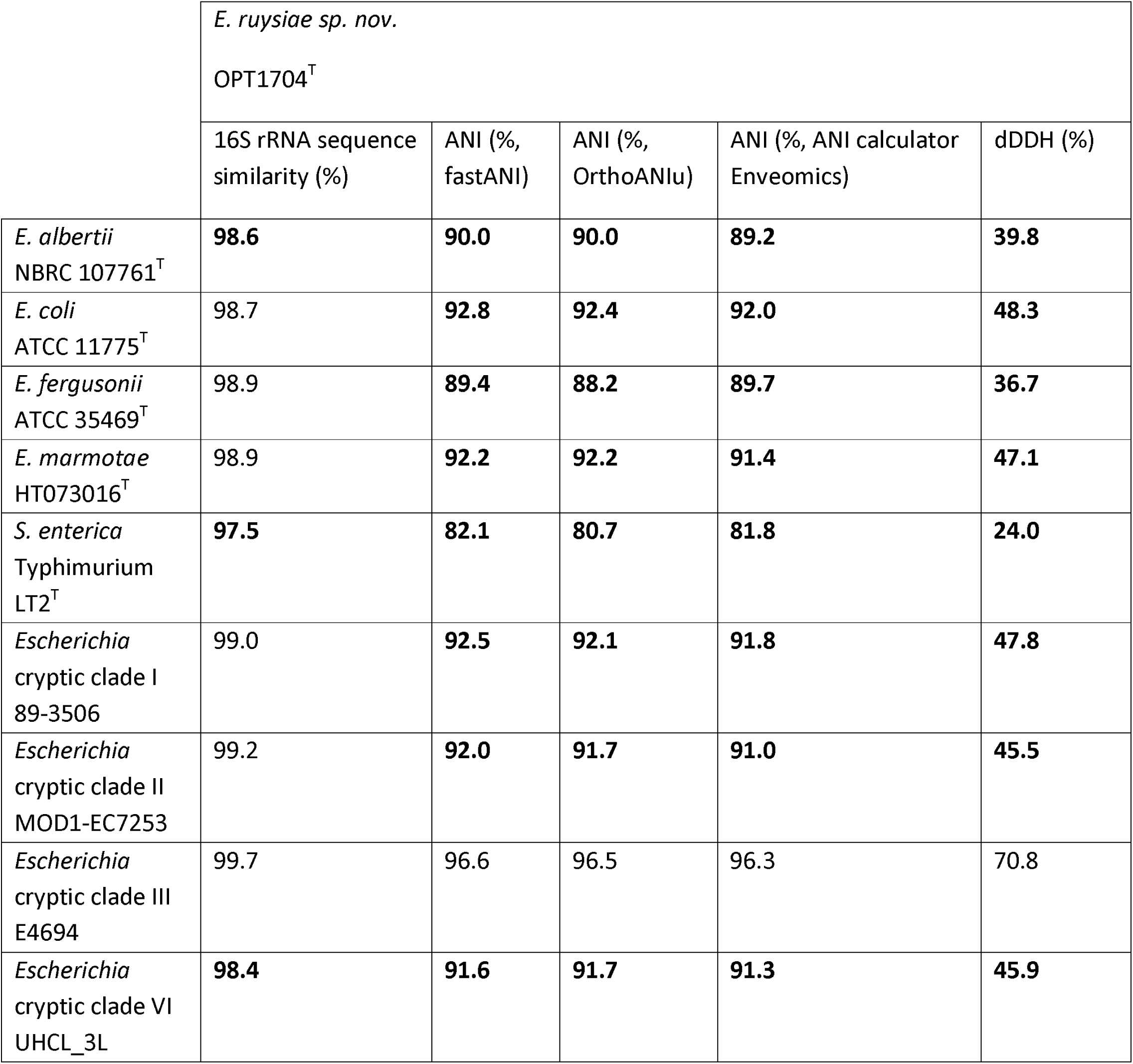
Comparison of OPT1704^T^ 16S rRNA and whole-genome sequence with type strains of *E. albertii, E. coli, E. fergusonii, E. marmotae*, representative genomes of *Escherichia* cryptic clades I, II, III and VI and *S. enterica* serovar Typhimurium. In bold are the values that warrant assignment of OPT1704^T^ to a novel species (<98.7% 16S rRNA sequence similarity, <95-96% ANI, <70% dDDH). ANI: average nucleotide identity, dDDH: digital DNA:DNA hybridisation.

|  | *E. ruysiae* sp. nov. |  |  |  |  |
| --- | --- | --- | --- | --- | --- |
|  | OPT1704<sup>T</sup> |  |  |  |  |
|  | 16S rRNA sequence similarity (%) | ANI (%<br>fastANI) | ANI (%<br>OrthoANlu) | ANI (%<br>ANI calculator<br>Enveomics) | dDDH (%) |
| *E. albertii*<br>NBRC 107761<sup>T</sup> | **98.6** | **90.0** | **90.0** | **89.2** | **39.8** |
| *E. coli*<br>ATCC 11775<sup>T</sup> | 98.7 | **92.8** | **92.4** | **92.0** | **48.3** |
| *E. fergusonii*<br>ATCC 35469<sup>T</sup> | 98.9 | **89.4** | **88.2** | **89.7** | **36.7** |
| *E. marmotae*<br>HT073016<sup>T</sup> | 98.9 | **92.2** | **92.2** | **91.4** | **47.1** |
| *S. enterica*<br>Typhimurium<br>LT2<sup>T</sup> | **97.5** | **82.1** | **80.7** | **81.8** | **24.0** |
| *Escherichia*<br>cryptic clade I<br>89-3506 | 99.0 | **92.5** | **92.1** | **91.8** | **47.8** |
| *Escherichia*<br>cryptic clade II<br>MOD1-EC7253 | 99.2 | **92.0** | **91.7** | **91.0** | **45.5** |
| *Escherichia*<br>cryptic clade III<br>E4694 | 99.7 | 96.6 | 96.5 | 96.3 | 70.8 |
| *Escherichia*<br>cryptic clade VI<br>UHCL_3L | **98.4** | **91.6** | **91.7** | **91.3** | **45.9** |

OPT1704^T^ showed 98.7-98.9% 16S rRNA sequence similarity to *E. coli* ATCC 11775^T^, *E. fergusonii* ATCC 35469^T^ and *E. marmotae* HT073016^T^, which would not warrant assignment to a novel species based on the current threshold for species delineation (less than 98.7% sequence similarity). However, the threshold for species delineation on the basis of 16S rRNA sequence has changed often and thresholds of up to 99% sequence similarity have been proposed previously (Kim 2014). In contrast, ANI analysis and dDDH did support assignment of OPT1704^T^ to a novel species, together with the representative strain of *Escherichia* cryptic clade III (table 2). The analyses also confirmed that OPT1704^T^ falls within the *Escherichia* genus. This novel species, encompassing both *Escherichia* cryptic clades III and IV, was assigned *E. ruysiae* sp. nov. with OPT1704^T^ as the proposed type strain.

To gain a better understanding of the *Escherichia* genus, we produced two phylogenies, based on 16S rRNA sequence (Fig. 1) and on an alignment of 682 core genes (Fig. 2). In short, rRNA genes were predicted from whole genomes using barrnap (version 0.9, https://github.com/tseemann/barrnap) and a tree was generated using FastTree (version 2.1.10 (26)). For the core gene alignment, genomes were first annotated with Prokka (version 1.14.0 (27)) and a core gene alignment was produced using Roary (version 3.12.0 (28)) and MAFFT (version 7.307 (29)). The phylogeny was inferred using a generalised time reversible model using base frequencies from the SNP alignment and free rate heterogeneity (GTR+F+R4 model) in IQ-tree (version 1.6.6 (30)), as advised by ModelFinder (31). Phylogenies were rooted on the *Salmonella enterica* serovar Typhimurium str. LT2^T^ genome. Both phylogenies showed that strain OPT1704^T^ clusters closely with the strain MOD1-EC7259 from *Escherichia* cryptic clade III, and away from the current *Escherichia* species.

**Figure 1.**
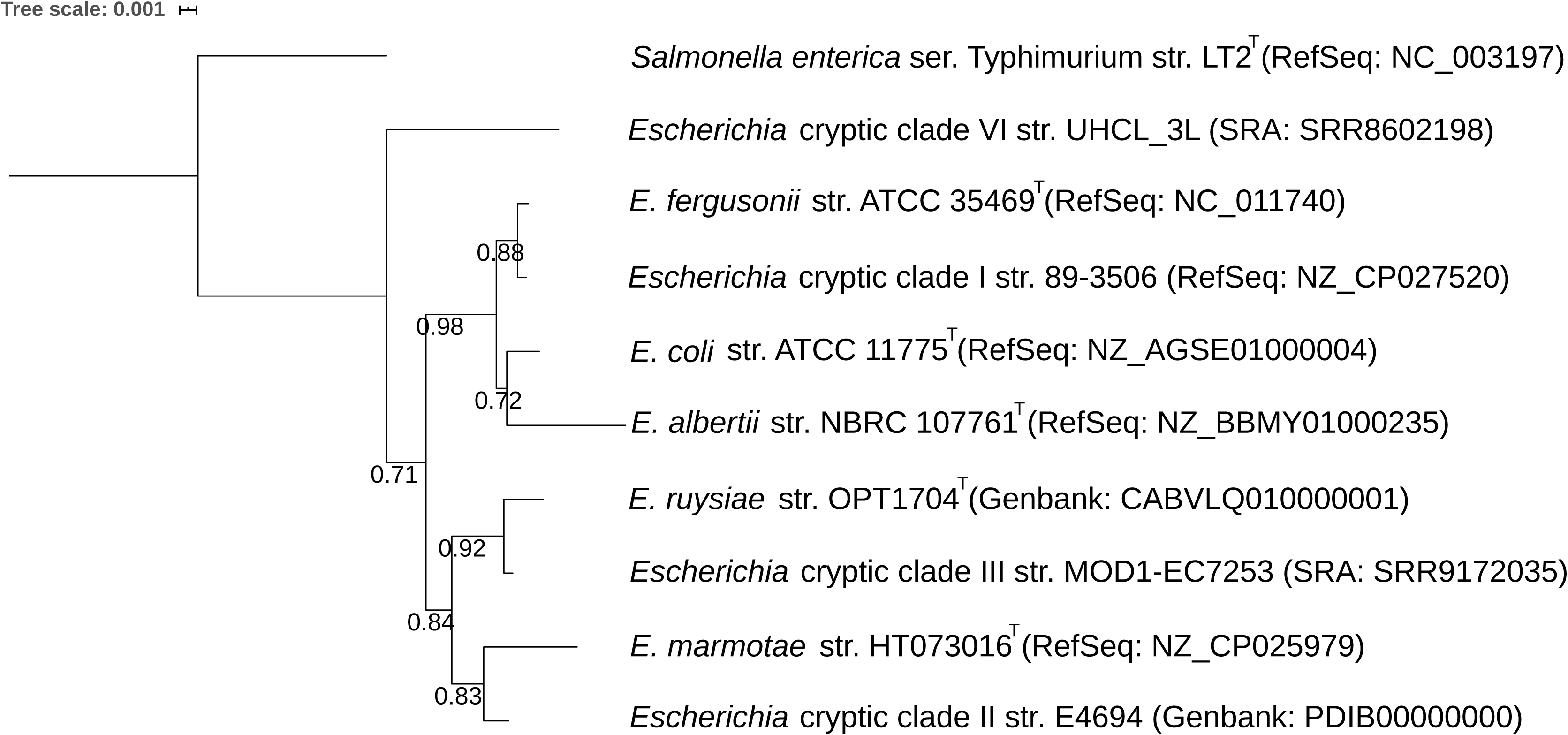
16S phylogeny of *E. ruysiae* str. OPT1704^T^ with type strains of other *Escherichia spp*., other *Escherichia* cryptic clades and *Salmonella enterica* serovar Typhimurium as outgroup. Numbers indicate bootstraps on a scale of 0 to 1. Phylogeny available at https://itol.embl.de/tree/14511722611197561579189697

**Figure 2.**
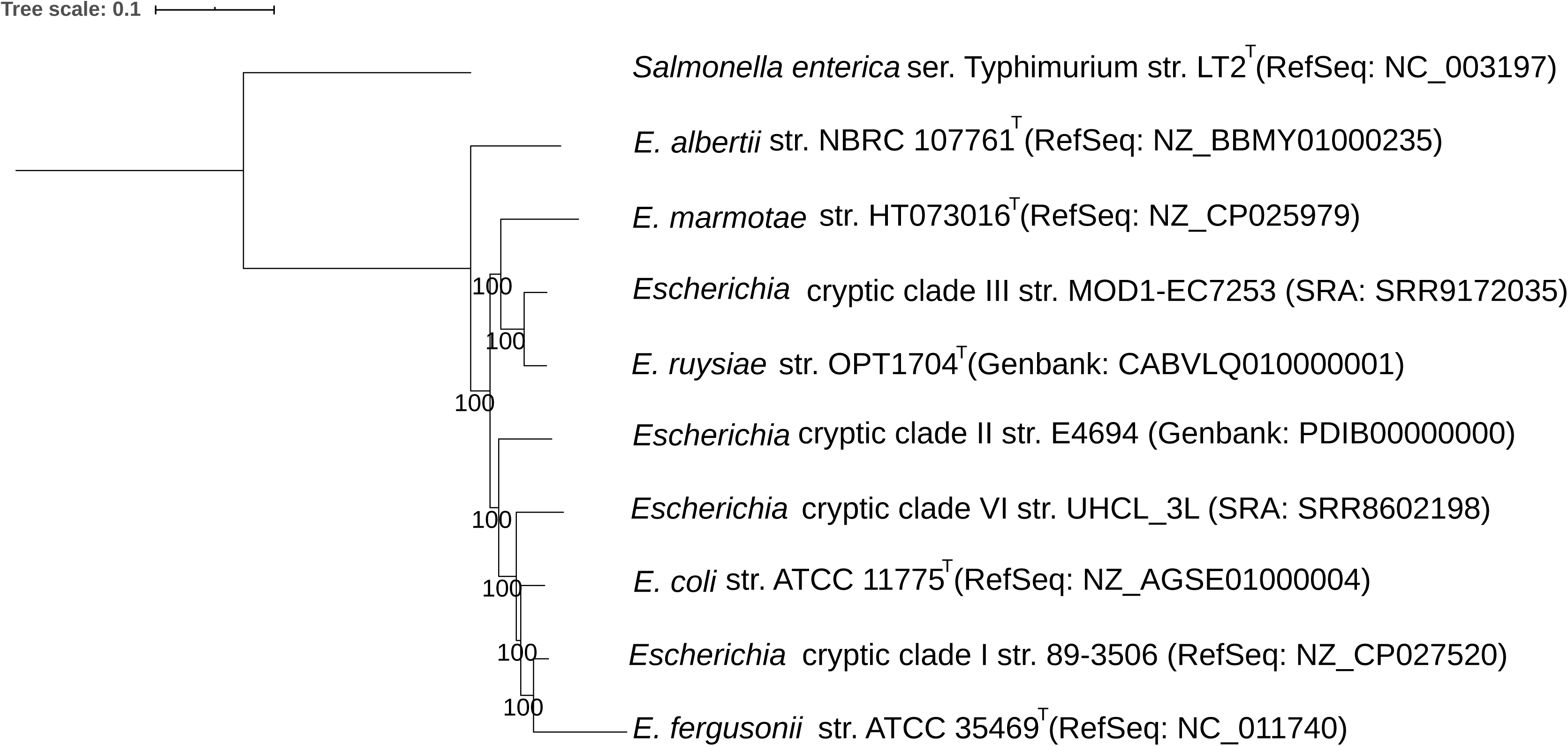
Phylogeny based on 682 concatenated core genes including *E. ruysiae* OPT1704T with type strains of other *Escherichia spp*., other *Escherichia* cryptic clades and *Salmonella enterica* serovar Typhimurium as outgroup. Numbers indicate bootstraps on a scale of 0 to 100. Phylogeny available at https://itol.embl.de/tree/14511722711358151579253758

Chun et al. (32) proposed that strains with >95-96% genome-wide ANI between each other should be assigned to the same species. If cryptic clades III and clade IV would share >95-96% ANI, this would mean both clades should be assigned to the same novel species, *E. ruysiae*. To assess this for a larger number of strains than the type strains presented in table 2, we downloaded all available WGS from clade III and clade IV strains from Enterobase and compared ANI between all genomes using fastANI (version 1.1 (22)). This analysis revealed that within 32 clade III genomes, the median ANI is 98.6% (range: 97.7%-99.9%), while within 31 clade IV genomes, the median ANI is 98.9% (range: 98.6%-99.9%). Between clade III and clade IV genomes, the median ANI is 96.6% (range 96.2%-96.8%). This suggests clades III and IV should be assigned to the same novel species, *E. ruysiae* sp. nov.

Currently, no IJSEM guidelines exist for the delineation of subspecies based on genomic data. However, *E. ruysiae* could potentially be delineated further into two subspecies (representing the current clades III and IV, respectively) in the future, after a type strain for cryptic clade III has been identified.

## Description of E. ruysiae sp. nov.

*Escherichia ruysiae* (ruy’si.ae N.L. fem. n. after Anna Charlotte Ruys, professor of microbiology at the University of Amsterdam from 1940 to 1969). Cells are Gram-negative, facultatively anaerobic, non-sporulating, non-motile rods with a size of approximately 1 by 2 µm. Colonies are circular, convex, grey-white and semi-transparent when grown overnight at 37 °C on COS sheep blood agar plates. The species is catalase-positive and oxidase-negative and grows at temperatures between 20 and 42 °C and NaCl concentrations between 0% and 6% w/v. In the VITEK2 GN biochemical test set it yields a positive result for Beta-Galactosidase, D-Glucose, D-Maltose, D-Mannitol, D-Mannose, D-Sorbitol, D-Trehalose, Saccharose/Sucrose, D-Tagatose, Gamma-Glutamyl-Transferase, Fermentation Glucose, Tyrosine Arylamidase, Succinate Alkalinisation, Alpha-Galactosidase, Ornithine Decarboxylase, Courmarate, Beta-Glucoronidase, 0/129 Resistance (Comp.Vibrio.) and Ellman and negative for Ala-Phe-Pro-Arylamidase, Adonitol, L-Pyrrolydonyl-Arylamidase, L-Arabitol, D-Cellobiose, H_2_S Production, Beta-N-Acetyl Glucosaminidase, Glutamyl Arylamidase Pna, Beta-Glucosidase, Beta-Xylosidase, Beta-Alanine Arylamidase Pna, L-Proline Arylamidase, Lipase, Palatinose, Urease, Citrate (Sodium), Malonate, 5-Keto-D-Gluconate, L-Lactate Alkalinisation, Alpha-Glucosidase, Beta-N-Acetyl-Galactosaminidase, Phosphatase, Glycine Arylamidase, Lysine Decarboxylase, L-Histidine Assimilation, Glu-Gly-Arg-Arylamidase, L-Malate Assimilation and L-Lactate Assimilation.

The 16S rRNA sequence is deposited in ENA under accession LR745848. Raw Illumina and Nanopore whole-genome sequencing data, as well as the complete genome assembly are deposited under project PRJEB34275.

## Acknowledgements

The authors would like to thank Rob Weijts and Patricia Brinke for their help in phenotypic characterization of type strain OPT1704^T^ of *Escherichia ruysiae* sp. nov. and Arie van der Ende for the helpful discussions. We thank SURFsara (www.surfsara.nl) for the support in using the Lisa Compute Cluster.

## AUTHOR STATEMENTS

### Ethical statement

Not required.

## ABBREVIATIONS

ONT: Oxford Nanopore Technologies
MIC: Minimum Inhibitory Concentration
COS: Columbia agar + Sheep blood
dDDH: digital DNA:DNA hybridisation
ANI: Average Nucleotide Identity
SNP: Single Nucleotide Polymorphism

